# The *E. coli* helicase does not use ATP during replication

**DOI:** 10.1101/2021.07.07.451541

**Authors:** Lisanne M. Spenkelink, Richard R. Spinks, Slobodan Jergic, Jacob S. Lewis, Nicholas E. Dixon, Antoine M. van Oijen

**Author notes:** These authors contributed equally to this work (Lisanne M. Spenkelink, Richard R. Spinks). Present Address: The Francis Crick Institute, 1 Midland Road, London NW1 1AT, U.K.

## Abstract

The replisome is responsible for replication of DNA in all domains of life, with several of its individual enzyme components relying on hydrolysis of nucleoside triphosphates to provide energy for replisome function. Half a century of biochemical studies have demonstrated a dependence on ATP as an energy source for helicases to unwind duplex DNA during replication. Through single-molecule visualization of DNA replication by the *Escherichia coli* replisome, we demonstrate that the DnaB helicase does not rely on hydrolysis of ATP (or any ribo-NTPs) in the context of the elongating replisome. We establish that nucleotide incorporation by the leading-strand polymerase is the main motor driving the replication process.

**One Sentence Summary:** Polymerases provide the energy for helicase-mediated DNA unwinding during *E. coli* DNA replication.

## Main Text

Adenosine triphosphate (ATP) hydrolysis is the main cellular source of energy to drive biochemical reactions that are otherwise energetically unfavorable. The chemical energy stored in phosphoanhydride bonds is released upon hydrolysis of ATP to ADP and is used to drive mechanical work and conformational change. A canonical process that is thought to rely on ATP hydrolysis is DNA replication, with the multi-enzyme replisome using the energy from ATP to promote activities such as DNA unwinding. A model system for DNA replication is the *Escherichia coli* replisome, which requires participation of 12 different proteins to duplicate the chromosome (Fig. 1A (inset)) (*1*). Of these, three components – the DnaB helicase, the DnaG primase and the clamp-loader complex – are known to use ATP in their function.

**Fig. 1.**
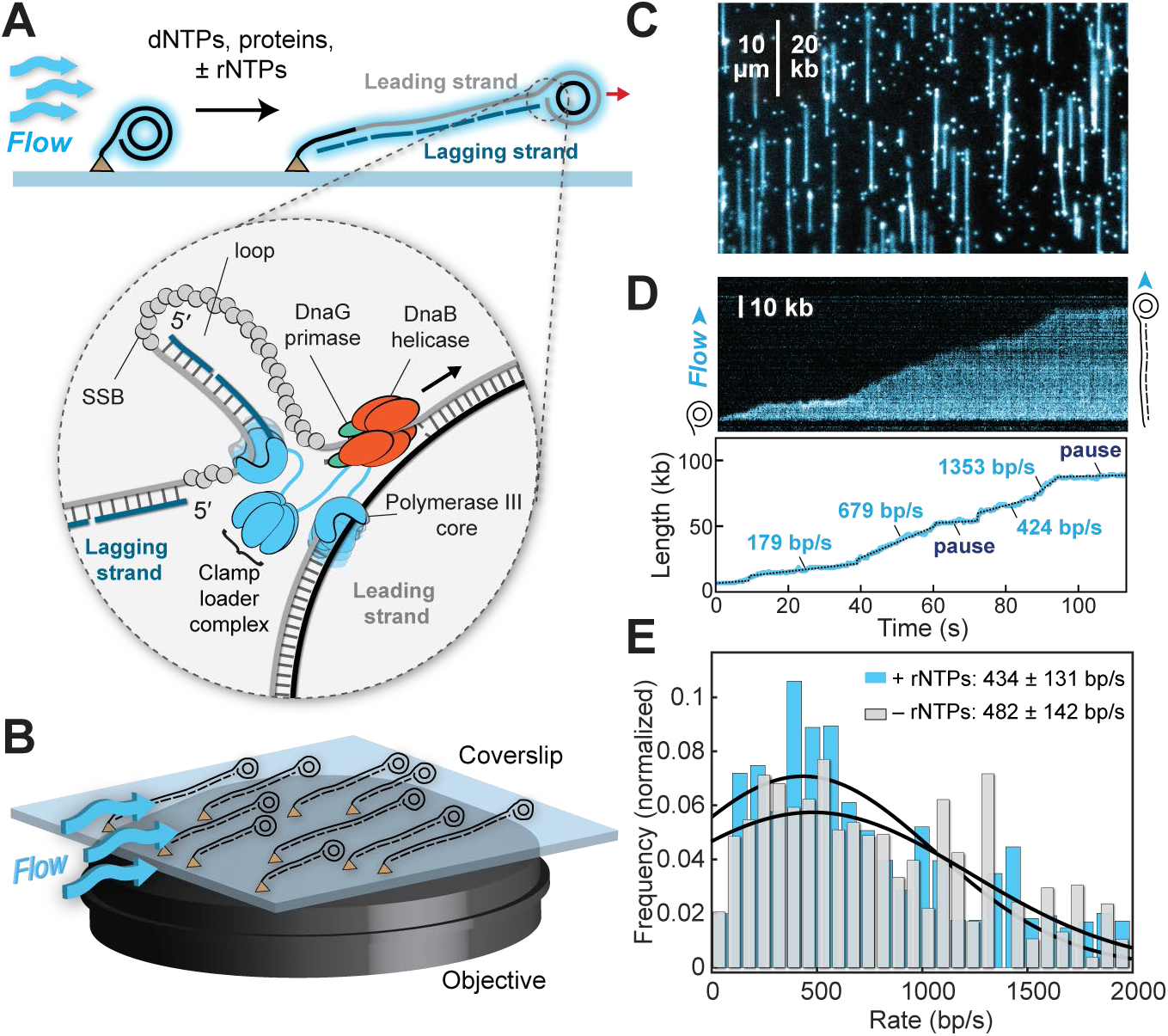
Single-molecule rolling-circle DNA-replication assay. (**A**) Schematic of the rolling-circle assay. Long products of leading- and lagging-strand DNA are stretched by the laminar flow of buffer. Inset: schematic representation of the *E. coli* replisome. (**B**) Rolling-circle DNA templates are immobilized on the surface of a microfluidic flow cell mounted on the objective of a TIRF microscope. (**C**) A typical field of view showing rolling-circle replication products. Scale bar = 10 μm. (**D**) Kymograph of an individual leading- and lagging-strand replication event. The position of the tip of the DNA corresponds the position of the replisome and the replication rate can be tracked over time using an automated tracking algorithm (blue). Individual rate segments are identified through change-point analysis (black). (**E**) Distributions of replication rates in the presence (blue, *N* = 58 molecules) and absence (gray, *N* = 58 molecules) of rNTPs, with Gaussian fit.

The replicative helicase DnaB hydrolyzes ATP or other ribonucleoside triphosphates, (rNTPs) (*2–7*) through its highly-conserved RecA-type ATPase domain (*8–11*). The energy from hydrolysis powers DnaB translocation along DNA (*3, 4, 7, 12–18*) to unwind the duplex at the replication fork (*19, 20*). The DnaG primase uses ATP as part of the *de novo* synthesis of RNA primers on the lagging strand (*21–25*). ATP forms part of the rNTP pool utilized by DnaG to generate RNA primers (*23*). The clamp-loader complex uses ATP to load the β_2_ processivity clamp onto DNA (reviewed in (*27*)).

Although these replisomal enzymes can use ATP in their function, not all of them completely depend on it. DnaG is a promiscuous primase and is capable of incorporating deoxy-NTPs (dNTPs) (*21, 23*). Similarly, while the clamp-loader complex predominantly uses ATP, it also accepts dATP as a substitute (*28, 29*). In contrast, the DnaB helicase is critically reliant on rNTPs. While DnaB can consume any of the four rNTPs, it has negligible capacity for dNTP hydrolysis to fuel translocation (fig. S1) (*2–5*).

The strict dependence of DnaB on rNTPs has led to the widely accepted view that DNA replication progresses in an ATP-dependent manner, a theory supported by ensemble biochemical studies that show a lack of DNA replication in the absence of ATP. However, most ensemble assays do not separate helicase loading from replisome activity and thus do not separately test the ATP dependence of replication elongation (unwinding and synthesis). Here we show, using three different single-molecule replication assays, that efficient replisome activity does not require ATP or other rNTPs. Our data suggest that the helicase can act passively, with the energy required for DNA-strand separation provided by the leading-strand DNA polymerase incorporating dNMPs.

Using a flow-cell based single-molecule rolling-circle replication assay, we are able to separate helicase loading and replication elongation into discrete steps and test the dependence of replisome-mediated elongation on ATP (*30–34*). First, with ATP present, we assembled DnaB from the DnaBC helicase-loader complex onto the rolling-circle DNA template, a 2-kb circular dsDNA template with a 5′ flap that mimics the forked DNA found at the site of replication (*35*). The helicase-DNA complex was immobilized on the surface of a microfluidic flow cell. Omitting ATP from this DnaB loading step resulted in a complete lack of replication, confirming the necessity of ATP during loading (fig. S2). Following helicase loading, and an extensive wash step to remove ATP, replication elongation was initiated by introducing the other replication components necessary for leading- and lagging-strand synthesis (see methods).

Leading-strand synthesis displaces ssDNA from the circle, with the liberated ssDNA subsequently acting as a template for lagging-strand synthesis (Fig. 1A). Thus, the rolling-circle design reconstitutes coupled leading- and lagging-strand synthesis and results in a single continuously growing dsDNA product. We stretch these growing DNA molecules by laminar flow and visualize a large number of them simultaneously by real-time near-TIRF wide-field imaging, using a dsDNA intercalator as a fluorescent probe (Fig. 1C; see methods). Using automated, unbiased tracking and change-point fitting algorithms (Fig. 1D) (*36–38*), we quantified the instantaneous rates of replication. In the presence of ATP, we find a replication rate of 434 ± 131 bp/s (mean ± s.e.m.), consistent with previous single-molecule observations (Fig. 1E) (*31, 32, 39*). Surprisingly, when all rNTPs are omitted from the replication elongation phase, we still detect efficient DNA replication with a similar instantaneous rate of 482 ± 142 bp/s (mean ± s.e.m.; Fig. 1E).

In the absence of rNTPs, we observed sporadic SSB-coated gaps in the lagging-strand product (fig. S3), which we attribute to inefficient Okazaki fragment priming by DnaG (*40*) as it incorporates dNTPs less efficiently than rNTPs (*21–23*). Notwithstanding these changes in primase behavior, the key observation is that replisome elongation rates are unaffected by the absence of rNTPs.

To more closely interrogate the replisomal mechanisms that allow the DnaB helicase to function without ATP, we simplified the reaction to observe only leading-strand synthesis (the components essential for lagging-strand activity were omitted; see methods) (Fig. 2A). Since this assay produces ssDNA instead of dsDNA, replication is visualized through imaging of ssDNA-bound fluorescently labeled single-stranded DNA-binding protein, SSB (Fig. 2B) (*33, 41*). In the presence or absence of ATP, we measure instantaneous replication rates of 420 ± 64 and 440 ± 60 nt/s, respectively, with similar replication efficiency (Fig. 2C, fig. S4), consistent with previous single-molecule observations that were done in the presence of ATP (*30*).

**Fig. 2.**
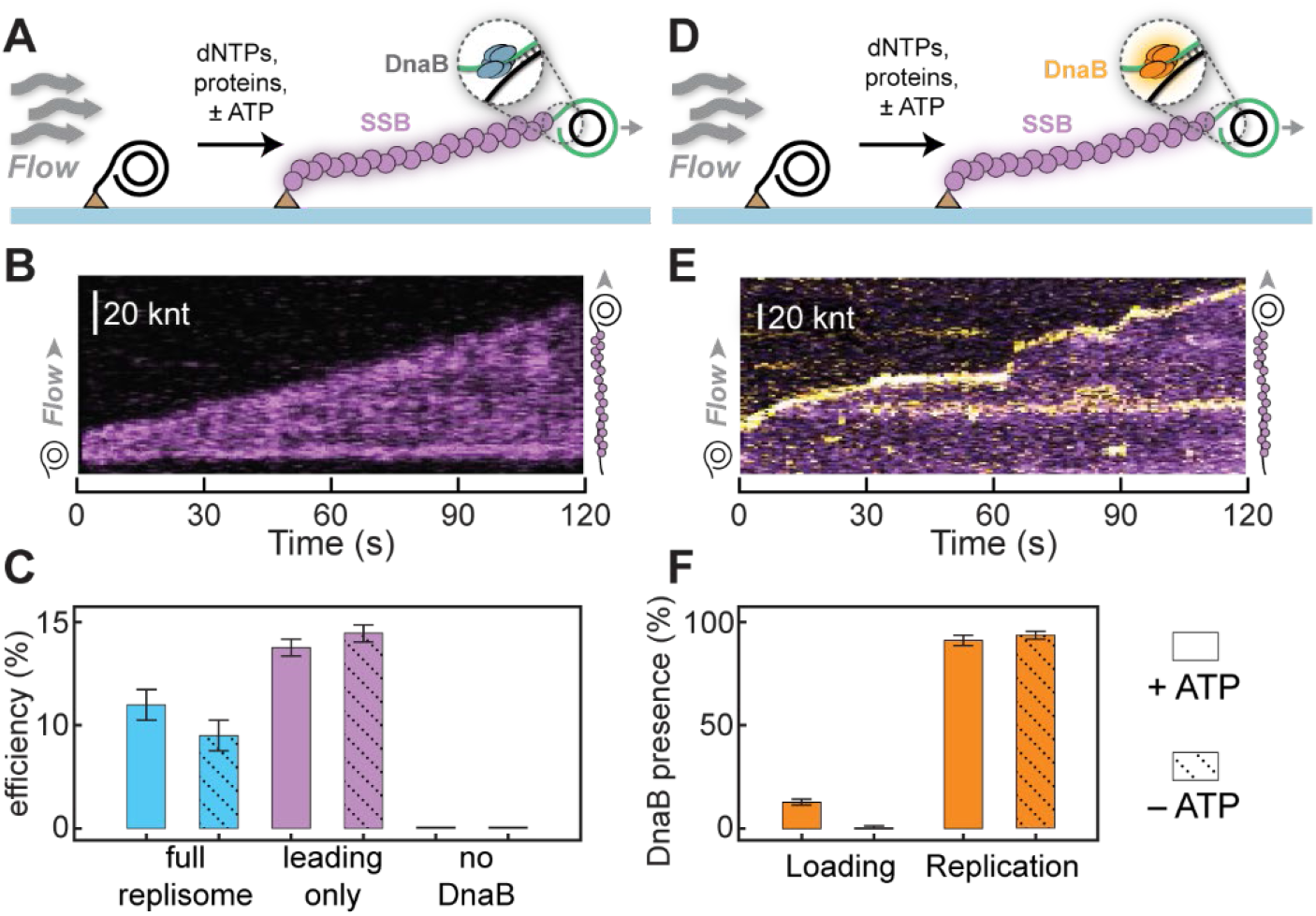
Leading-strand synthesis assay confirms the presence of DnaB irrespective of ATP. (**A**) The assay is set up as before, with rNTPs and DnaG primase omitted. ssDNA is visualized using fluorescently labeled SSB (purple). (**B**) Representative kymograph of leading-strand replication. Fluorescently labeled SSB stains the growing ssDNA product (green). (**C**) Replication efficiencies for leading- and lagging-strand synthesis (blue), leading-strand synthesis only (purple), and leading-strand synthesis in the absence of DnaB (black) ± ATP. (**D**) Fluorescent DnaB is added to monitor the presence of DnaB at the site of replication. (**E**) Representative kymograph showing the presence of DnaB (orange) at the tip of the ssDNA replication product (purple). (**F**) Quantification of the number of DNA products that have DnaB present at the fork in loading and in replication ± ATP. Data for DnaB loading is presented in fig. S7.

To confirm that the observed replication products are indeed the result of helicase-mediated synthesis, we carried out two important controls. First, the leading-strand synthesis assay was repeated with the simultaneous two-color visualization of fluorescent SSB and DnaB helicase fluorescently labeled in a different color (*34*) (Fig. 2D,E). We show fluorescent DnaB is loaded only when ATP is present (Fig 2F, fig. S2). However, once loaded we detect fluorescent DnaB in all elongating replisomes, irrespective of the presence of ATP during replication (Fig. 2F, fig. S7). For our second control, we repeated leading-strand synthesis without DnaB and found no replication products (Fig. 2C, fig. S5). From these two observations we reaffirm that DnaB is a necessary part of the replisome but does not require ATP to support replication elongation.

Next, we sought to determine which energy source, if not helicase-mediated ATP hydrolysis, allows the replisome to unwind DNA. SSB is known to have passive unwinding activity whereby it binds to ssDNA that is transiently exposed at a ssDNA-dsDNA fork due to thermal breathing of the dsDNA (*42, 43*). The bound SSB then potentially acts as a ratchet, preventing the reannealing of the ssDNA. Although SSB is not necessarily required for leading-strand synthesis in the presence of ATP (*30*), it is conceivable that SSB binding rather than helicase activity enables fork progression under conditions without ATP. To exclude this possibility, we designed a minimal replication assay in which we also omit SSB. We carried out the assay in the absence of flow (Fig. 3A) so that the exposed ssDNA product from rolling-circle leading-strand synthesis forms an entropic coil with high local concentration of ssDNA, which can be stained and visualized using the low-affinity binding of the DNA fluorescent stain to ssDNA. We monitor replication through an increase in intensity of the stained DNA product (Fig. 3B, C). The assay was calibrated using the intensity of a ssDNA template of known length (fig. S6). Using change-point analysis we find an instantaneous rate of replication of 368 ± 157 nt/s in the presence of ATP (fig. S8). This rate is comparable to our prior rate measurements with SSB (fig. S4) and to previously reported rates from single-molecule experiments in the absence of SSB (*30*). Again, we see efficient replication (340 ± 277 nt/s) in the absence of ATP (fig. S6). The similarity of these rates indicates that SSB is not responsible for strand separation.

**Fig. 3.**
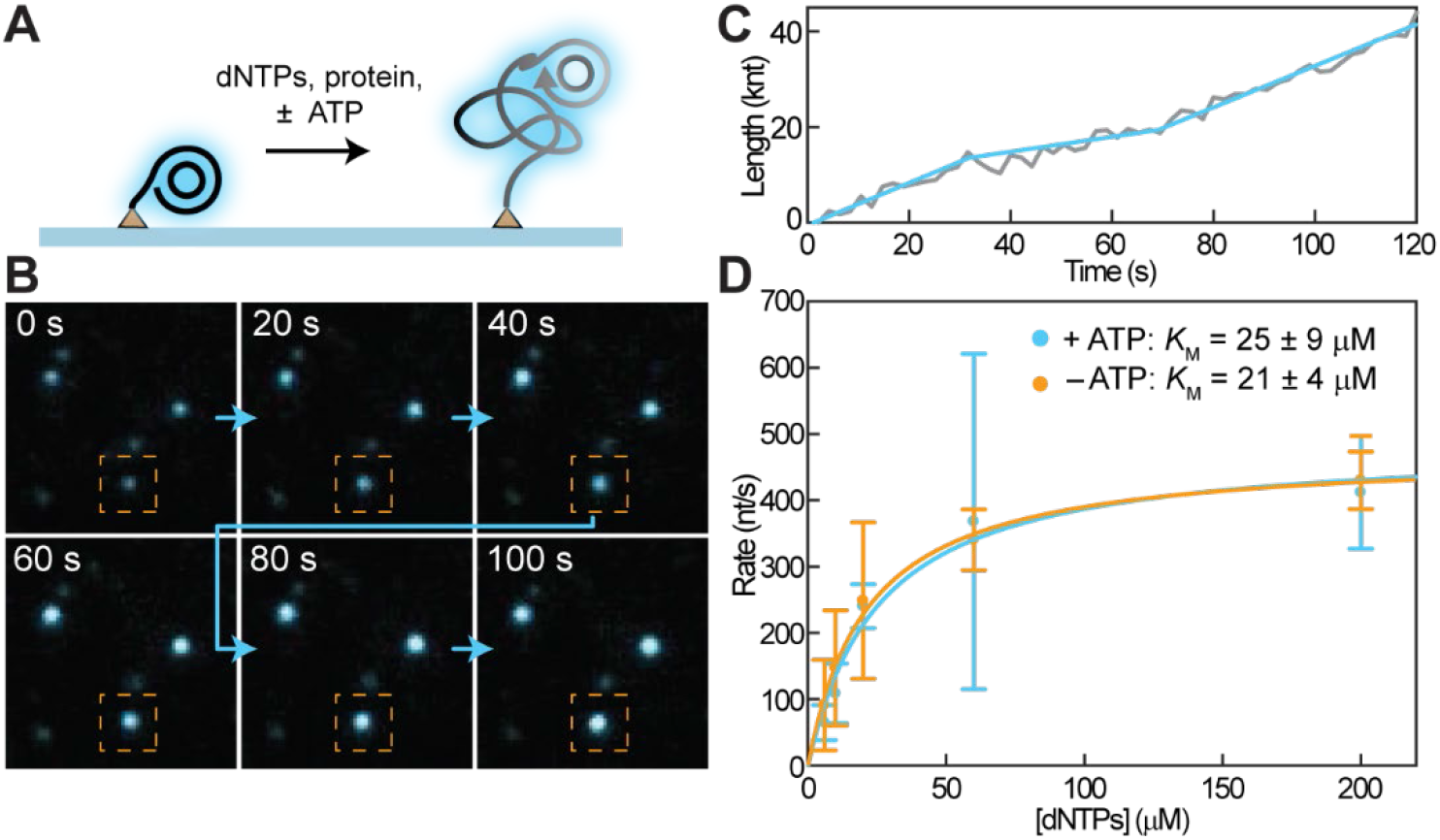
Minimal-replisome assay provides evidence that the polymerase is the main driving force of the replisome. (**A**) Schematic of the assay. In the absence of flow, the newly synthesized DNA forms a compact coil of ssDNA. (**B**) Montage showing the increase in size and intensity of three individual DNA molecules undergoing replication. (**C**) Intensity is converted into length and plotted as a function of time for the boxed molecule in B (gray), and rate segments determined by change-point fitting (blue). (**D**) Rate of replication as a function of dNTP concentration in the presence (blue) and absence (orange) of ATP; [dNTPs] represents the concentration of each of the four dNTPs. The solid lines represent Michaelis-Menten fits to the data.

Finally, we consider the possibility that DnaB might hydrolyze dNTPs when associated with the replisome. DnaB has previously been shown not to have detectable ssDNA-dependent dNTPase activity (*3*). The structural basis for this specificity is a H-bond to the 2′-OH of ATP, which would be absent in dNTPs (*10*). Using an assay that measures the rate of dissociation of DnaB from a ssDNA template with a blocked 3′ end as a proxy for its affinity for NTPs (fig. S1), we confirm that its preference for dATP is ∼1000-fold lower than for ATP. With the *K*_m_ of DnaB for ATP hydrolysis previously determined to be ∼100 μM (*3*), this observation implies a *K*_m_ for the dATPase activity of DnaB of about 100 mM. To assess whether DnaB might display an improved ability to hydrolyze dNTPs (*i.e*., have a much lower *K*_m_ for dNTPs) when associated with the replisome, we set out to measure replication rates as a function of dNTP concentration in the presence and absence of ATP. First, we determine the *K*_m_ of replication for dNTPs in the presence of ATP by measuring the rate of replication in our minimal replication assay as function of dNTP concentration in the presence of ATP in our minimal replication assay. As expected, the rate of replication increases with increasing concentrations of dNTPs. A fit of the data (Fig. 3D) gives a *K*_m_ (per dNTP) of 25 ± 9 μM. By leaving out ATP in the same reaction we will be able to measure the *K*_m_ of DnaB for dNTPs, as long as that *K*_m_ is higher than 25 μM. This lower limit would allow us to observe decreases in *K*_m_ of DnaB for dNTPs by up to 4,000 fold. In the absence of ATP we find a *K*_m_ of 21 ± 4 μM (Fig. 3D). These unchanged *K*_m_ values demonstrate that the rate limiting factor in DNA replication is utilization of dNTPs by the polymerase, not by the helicase. This observation means that when associated with the replisome, either the DnaB does not utilize dNTPs or that it has a *K*_m_ for dNTPs more than 4,000 times lower than when acting alone, an extremely unlikely scenario.

Taken together, our data suggest that neither hydrolysis of ATP nor any other r/dNTP by the helicase is absolutely required for DNA strand separation during replication. We propose that dNTP incorporation by the leading-strand polymerase is the main driving force for replisome progression. Notably, in the absence of DnaB, polymerase III holoenzyme coupled to SSB is also capable of unwinding dsDNA and DNA synthesis, albeit with much lower rates and with higher *K*_m_ for dNTPs (*44, 45*). This observation indicates the critical role that DnaB still plays during replication, even when not hydrolyzing rNTPs.

It has been shown previously using the simpler bacteriophage T7 replisome that the replicative polymerase and helicase each stimulates the activity of the other, indicating an active role of the polymerase in driving DNA replication at maximal rates (*46*). Moreover, a cryo-EM structure of the T7 replisome shows the polymerase positioned perpendicular to the helicase at the DNA fork junction, also implying a cooperative approach to DNA unwinding (*47*). Hence it seems likely that DnaB provides the platform that dictates the architecture of the *E. coli* replisome to facilitate strand separation. Finally, while we show that rNTPs are not required during unimpeded replication, it is tempting to speculate that the helicase could use the energy from ATP hydrolysis to sustain replication through secondary structures or during roadblock bypass.

## Funding

This work was supported by the Australian Research Council (grants DP180100858 and DP210100067 to A.M.v.O. and N.E.D.; and an Australian Laureate Fellowship FL140100027 to A.M.v.O.), and an Australian Government Research Training Program Scholarship (to R.R.S). Funding for open access charge: Australian Research Council.

## Author contributions

Conceptualization, L.M.S., R.R.S., J.S.L., N.E.D., A.M.v.O.; Methodology, L.M.S., R.R.S, S.J., J.S.L., N.E.D., A.M.v.O.; Software, L.M.S.; Validation, L.M.S., R.R.S.; Formal analysis, L.M.S., R.R.S.; Investigation, L.M.S., R.R.S., S.J., J.S.L.; Resources, R.R.S., S.J.; Funding acquisition, N.E.D., A.M.v.O.; Supervision, N.E.D., A.M.v.O.; Visualization, L.M.S., R.R.S., S.J.; Writing – original draft, L.M.S., R.R.S., S. J., N.E.D., A.M.v.O.

## Competing interests

Authors declare no competing interests.

## Data and materials availability

Data available upon request. Analysis software is available on github: https://github.com/SingleMolecule.

## Materials and Methods

### Proteins

*E. coli* DNA replication proteins were produced as described previously: the β_2_ sliding clamp (*48*), SSB (*49*), AF647- and AF488-labeled SSB-K43C (*33*), the DnaB_6_(DnaC)_6_ helicase–loader complex, DnaC loader and AF647-labeled DnaB_6_H201C (*34*), DnaG primase (*50*), the τ_3_δδ′_χψ_ clamp loader (*30*), and Pol III αεθ core (*32*,*45*).

### Single-molecule rolling-circle experimental design

Construction of the 2030-bp template used for most rolling-circle assays has been described (*35*). Microfluidic flow cells were prepared as described (*51*). Briefly, a PDMS flow chamber was placed on top of a PEG-biotin-functionalized microscope coverslip. To help prevent non-specific interactions of proteins and DNA with the surface, the chamber was blocked with buffer containing 50 mM Tris-HCl pH 7.5, 50 mM KCl, and 2% Tween-20. The chamber was placed on an inverted microscope (Nikon Eclipse Ti-E) with a CFI Apo TIRF 100x oil-immersion TIRF objective (NA 1.49, Nikon) and connected to a syringe pump (Adelab Scientific) for flow of buffer. Reactions were carried out at 31°C, maintained by an electrically heated chamber (Okolab). Double-stranded DNA was visualized in real time by staining it with 150 nM SYTOX Orange (Invitrogen) excited by a 514-nm laser (Coherent, Sapphire 514-150 CW) at 150 μW/cm^2^. The red-labeled SSB was excited at 700 μW/cm^2^ with a 647 nm laser (Coherent, Obis 647-100 CW) and the blue-labeled SSB was excited at 700 μW/cm^2^ with a 488-nm laser (Coherent, Sapphire 488-200 CW). The red-labeled DnaB was excited at 100 μW/cm^2^ with the 647-nm laser. Imaging was done with an EMCCD camera (Hamamatsu). Signals were separated *via* dichroic mirrors and appropriate filter sets (Chroma).

### Leading- and lagging-strand replication assay

Conditions for simultaneous leading- and lagging-strand DNA replication were adapted from previously described methods (32– *34*). Briefly, 4 nM DnaB_6_(DnaC)_6_ was pre-loaded onto 20 pM rolling-circle DNA template by incubation at 37°C for 30 s in replication buffer (25 mM Tris-HCl pH 7.9, 50 mM potassium glutamate, 10 mM Mg(OAc)_2_, 40 μg/ml bovine serum albumin, 0.1 mM EDTA and 5 mM dithiothreitol) + 1 mM ATP. This mixture was loaded into the flow cell at 70 μl/min for 60 s and then at 10 μl/min for 8 min. To remove any unbound DNA and ATP from solution, the flow cell was vigorously washed with 100 flow-cell volumes of replication buffer. The reaction buffer consisted of 60 μM of each dNTP, 1.25 mM ATP and 250 μM each of CTP, GTP and UTP (when indicated) in replication buffer. Pol III* was assembled *in situ* by incubating τ_3_δδ′_χψ_ (410 nM) and Pol III cores (1.2 μM) in reaction buffer at 37°C for 90 s. Replication was initiated by flowing in the reaction buffer containing 3 nM Pol III*, 30 nM β_2_, 75 nM DnaG, and 20 nM SSB_4_ at 10 μl/min.

### Leading-strand replication assay

DnaB_6_(DnaC)_6_ was pre-loaded on the rolling-circle DNA template before immobilization of the helicase–DNA complex on the surface of the microfluidic flow cell, as described above. To remove any unbound template and ATP from solution, the flow cell was washed with 100 flow-cell volumes of replication buffer. The reaction buffer contained 60 μM of each dNTP and 1 mM ATP as indicated. Pol III* was assembled *in situ* by incubating τ_3_δδ′_χψ_ (410 nM) and Pol III cores (1.2 μM) in replication buffer at 37°C for 90 s. Replication was initiated by flowing in the reaction buffer containing 3 nM Pol III*, 30 nM β_2_, and 20 nM labeled SSB_4_ at 10 μl/min.

### Quantification of the length of SSB-coated ssDNA

We estimated the length of SSB-coated ssDNA in our assay based on the intensity of the labelled SSB. We divided the total intensity of 16 leading-strand synthesis products by the intensity of a single SSB. We assumed the SSB binds in the 35-mode to obtain a conversion factor of 752 ± 411 nt/pix (Fig S4A).

### DnaB loading quantification

4 nM labeled DnaB_6_(DnaC)_6_ was pre-loaded onto 20 pM rolling-circle DNA template as described above. This complex with 150 nM SYTOX orange was loaded into the flow cell at 70 μl/min for 60 s and then at 10 μl/min for 8 min. We imaged SYTOX stained DNA and labeled DnaB sequentially. We determined the position of DNA and DnaB foci using a peak-finding algorithm that fits 2D gaussian to give sub-pixel localisation. Foci were defined as colocalized when they were within 2 pix of eachother. Colocalisation by chance was calculated bases on area overlap (*32*).

### Minimal replication assay

DnaB_6_(DnaC)_6_ was pre-loaded on the rolling-circle template before immobilization of the helicase–DNA complex on the surface of the microfluidic flow cell, as described above. To remove any unbound template and ATP from solution, the flow cell was washed with 100 flow-cell volumes of replication buffer. The reaction buffer contained 60 μM of each dNTP and 1 mM ATP as indicated. Pol III* was assembled *in situ* by incubating τ_3_δδ′_χψ_ (410 nM) and Pol III cores (1.2 μM) in replication buffer at 37°C for 90 s. Replication was initiated by flowing in the reaction buffer containing 3 nM Pol III*, 30 nM β_2_, and 150 nM SYTOX orange at 10 μl/min.

### Calibration of SYTOX intensity as a function of ssDNA length

To quantify the rates of replication in the minimal assay, the intensity of SYTOX stained ssDNA was calibrated using M13 ssDNA (fig S6). A 66-mer 5′-biotin-T36AATTCGTAA TCATGGTCATAGCTGTTTCCT-3′ (Integrated DNA Technologies) was annealed to M13mp18 ssDNA (Guild Biosciences). This ssDNA template was loaded on the flow cell in reaction buffer and imaged using conditions identical to those used during the minimal replication assay. Using the average intensity measured in this assay and the known length of M13mp18 (7429 nt), an intensity per nt conversion factor can be obtained.

### Quantification and statistical analysis of single-molecule experiments

All analysis was done with ImageJ and MATLAB using in-house built plugins. The position of the tip of growing rolling-circle products was tracked as a function of time using an automated tracking algorithm. Individual rate segments were identified using an unbiased change-point algorithm (*36 – 38*). In the minimal replication assay, single-molecule trajectories were obtained by tracking the intensity of the SYTOX stained ssDNA product over time. The change of intensity in time (rate) was converted to nt/s using the ssM13 calibration. The rate histograms were weighted by the length of the segments to reflect the higher confidence for longer segments.

All single-molecule experiments were carried out in triplicate. The number of molecules or events analyzed is indicated in the text or figure legends. Errors reported in this study represent the standard error of the mean (s.e.m.) or the error of the fit, as indicated in the text or figure legends.

### Nucleotide-dependent DnaB dissociation assay using surface plasmon resonance (SPR)

SPR experiments used a BIAcore T200 (Cytiva) instrument at 25°C. First, a streptavidin-coated (SA) sensor chip (Cytiva) was activated with three sequential injections of 1 M NaCl, 50 mM NaOH (40 s each at 5 μl/min). Then, 50-mer ssDNA oligonucleotides were immobilized separately on two flow cells to 70 and 67 RU (response units), respectively, in SPR buffer (20 mM Tris pH 7.6, 150 mM NaCl, 100 μM EDTA and 0.005% *v/v* surfactant P20) supplemented with 0.5 mM ATP for subsequent stabilization of the DnaB helicase on the ssDNA.

Two biotinylated (bio) oligonucleotide sequences, exposing either a free 5′ or 3′ end were used: 5′-(T)_25_ GCA GGC TCG TTA CGT AGC TGT ACC G-bio-3′ (5′ EF-DNA) and 5′-bio-GCA GGC TCG TTA CGT AGC TGT ACC G(T)_25_-3′ (3′ EF-DNA).

In a standard experiment, DnaB was immobilized on the two separate ssDNA templates (5′ EF-DNA and 3′ EF-DNA) by injecting 250 nM DnaB_6_(DnaC)_6_ in SPR buffer supplemented with 1 mM ATP and an additional 10 mM EDTA to sequester excess Mg^2+^ present in the protein stock (note the absence of Mg^2+^ in the buffer) at a flow rate of 3 μl/min for 350 s. Subsequently, a ∼3,000 s wash step at a flow rate of 20 μl/min in SPR buffer + 0.5 mM ATP (running buffer) was applied to allow dissociation of DnaC from DnaB. From here, nucleotide-dependent DnaB dissociation experiments could be carried out for the exposed 5′- and 3′- end ssDNA templates with a variety of nucleotides.

For each condition, SPR buffer + 5 mM MgCl_2_ was injected with a particular r/dNTP or ADP at 1 mM concentration, at a flow rate of 20 μl/min for 1000 s simultaneously over both flow cells. For the ATP-titration experiments, DnaB dissociation from the 5′ EF-DNA template was monitored in SPR buffer + 5 mM MgCl_2_ supplemented with 1, 5, 20, 100 or 1000 μM ATP. Between each round of experiments, the surface was first regenerated by two 40 s injections of 3 M MgCl_2_ at 10 μl/min, and DnaB was re-immobilized onto the surface as described above. During DnaC rebinding experiments, 1 μM DnaC in SPR buffer supplemented with 1 mM ATP and 5 mM EDTA was injected at 20 μl/min for 200 s (fig. S1).

## Supplementary Figures

**Fig. S1.**
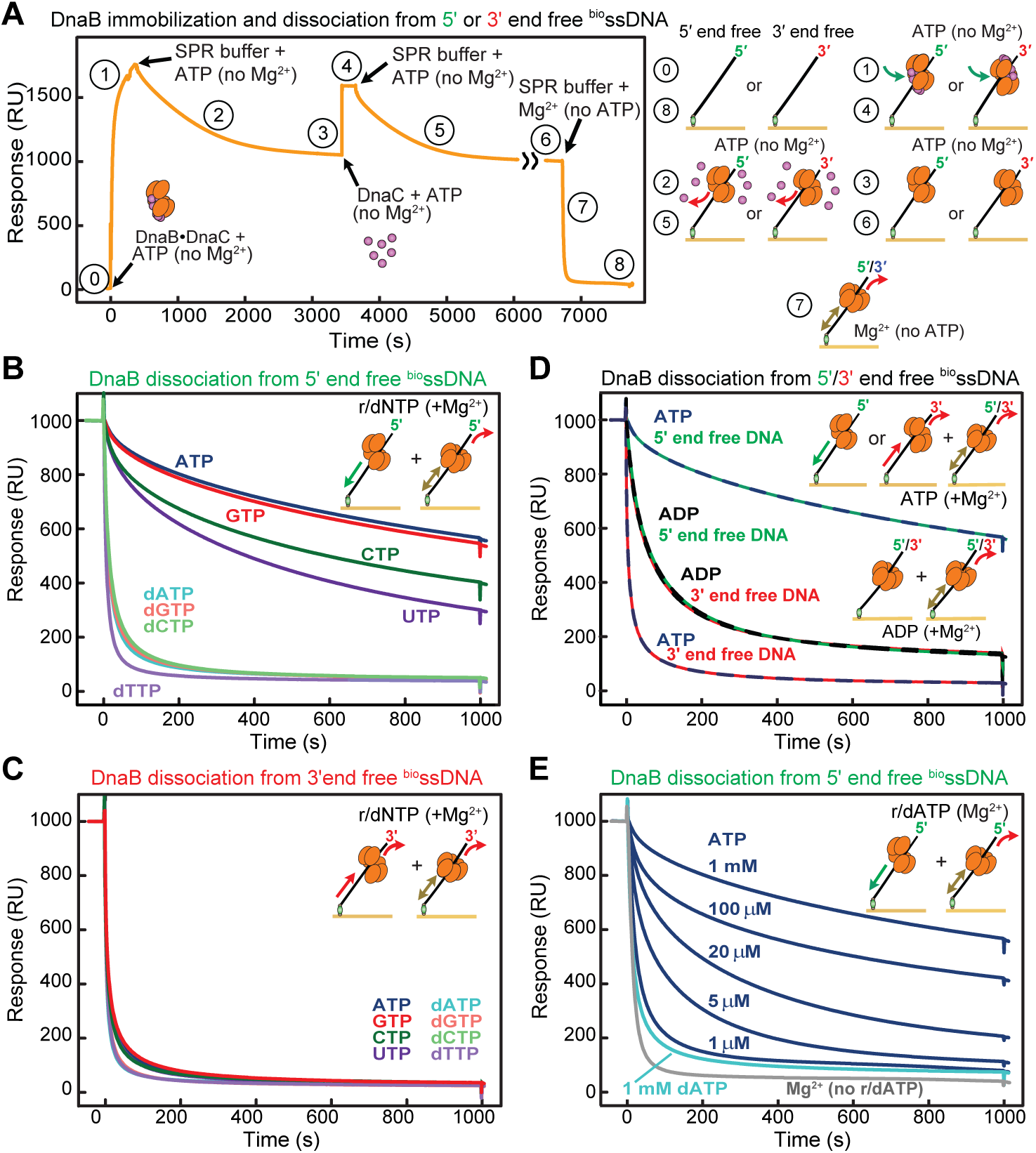
Comparison of nucleotide-dependent DnaB activities on short ssDNA templates using surface plasmon resonance (SPR). **(A)** SPR experimental design – stable immobilization of DnaB helicase on ssDNA and its nucleotide-dependent dissociation. We adapted methods from (*52*) to obtain stable association of hexameric DnaB on immobilized 50-mer ssDNA oligonucleotides with either an exposed 5′ or 3′ end (step 0); a representative SPR sensorgram is shown on the left, and pictorial representations of steps on the right. Stable association of DnaB_6_(DnaC)_6_ on ssDNA is achieved by omitting Mg^2+^ from the SPR buffer, thus permitting DnaB to bind ATP but not to hydrolyze it (step 1). Association is followed by a ∼3000 s wash step in the same buffer (step 2), in which DnaC is able to almost fully dissociate (step 3). We found immobilized DnaB to be very stably-bound in absence of Mg^2+^ and that DnaC could be re-associated (step 4), and then dissociated again (steps 5 and 6) in repeated cycles. Based on the difference in response units (RU) between DnaC cycles (step 4), we calculate that ∼90% of immobilized DnaB hexamers were retained over this period. Finally, with 1000 s of buffer injection with 5 mM Mg^2+^ but no ATP, we detected the relatively prompt dissociation of DnaB from ssDNA (steps 7 and 8). Herein lies the basis of this assay: the retention of DnaB can be compared on either exposed 5′- or 3′-end ssDNA in the presence of different nucleotides to determine the effects of these nucleotides on ssDNA binding and/or translocation. Note that translocation on the exposed 5′-end DNA would leave DnaB blocked at the chip surface, while its translocation on the exposed 3′-end DNA would result in the prompt dissociation of the helicase. For direct comparison, all signals were normalized to 1000 RU prior to induced DnaB dissociation. Injection spikes were also removed from the SPR sensorgrams. **(B)** rNTPs more efficiently retain DnaB on exposed 5′-end ssDNA over dNTPs. Real-time dissociation of DnaB from 5′-end ssDNA in the presence of 1 mM r/dNTP (as indicated) in SPR buffer with 5 mM Mg^2+^. The measurements (a) confirm the strong preference of rNTP over dNTP to stabilize the DnaB on ssDNA, either due to the static stabilization of the interaction with DNA or its active translocation in the 5′ − 3′ direction whereby the DnaB is likely to remain topologically trapped on DNA, and (b) demonstrate that of the four rNTPs, DnaB displays clear preference for purine rNTPs (ATP over GTP) over pyrimidines (CTP over UTP) for DNA binding and/or translocation. These trends mirror its preferences among rNTPs in its ssDNA-dependent rNTPase activities (*3*). **(C)** Neither rNTPs nor dNTPs efficiently retain DnaB on exposed 3′-end ssDNA. Real-time dissociation of DnaB from the 3′-end ssDNA in the presence of 1 mM r/dNTP (as indicated) in SPR buffer with 5 mM Mg^2+^. The results show fast dissociation of DnaB from the 3′-end ssDNA with all nucleotides. With dNTPs, DnaB likely has less affinity for ssDNA permitting the quick release of DNA and dissociation *via* diffusion. For rNTPs, the much faster dissociation of DnaB from the 3′-end ssDNA compared to 5′-end ssDNA (compare panels B and C) underscores the role of rNTP hydrolysis in active DnaB translocation. **(D)** Observation of ATP-hydrolysis driven translocation of DnaB on ssDNA. Real-time dissociation of DnaB from exposed 3′- and 5′-end ssDNA templates were compared in the presence of either 1 mM ATP or ADP and 5 mM Mg^2+^. At 1 mM ADP, DnaB is somewhat retained on both DNA templates (overlapping black/green and black/red sensorgrams, respectively) because ADP can only promote static binding of DnaB to ssDNA. In contrast, ATP affects the stability of DnaB on two different DNA templates antagonistically. On both templates, DnaB will use ATP hydrolysis to stimulate translocation in the 5′–3′ direction. On the 5′-end ssDNA, translocation topologically traps DnaB at the surface, but on the 3′-end ssDNA, translocation accelerates the dissociation of DnaB from the template. **(E)** DnaB utilizes rATP at least 1000-fold more efficiently than dATP. Real-time dissociation of DnaB from exposed 5′-end ssDNA was compared in SPR buffer with 5 mM Mg^2+^ as ATP was titrated in a the 1–1000 μM range (including zero) or with 1 mM dATP. Previous studies based on measurements of ATP hydrolysis have found DnaB-ssDNA has a *K*_m_ value of 106 μM for ATP (*3*). A somewhat lower *K*_m_ for ATP could be inferred from our titration profile, presumably due to the population of DnaB stabilized on ssDNA but not hydrolyzing ATP. Nevertheless, our results indicate >1000-fold higher propensity of DnaB helicase to utilize ATP over dATP (*i.e.*, DnaB still dissociates more slowly in 1 μM ATP than in 1 mM dATP), implying a >1000-fold lower *K*_m_ value for ATP over dATP in the process.

**Fig. S2.**
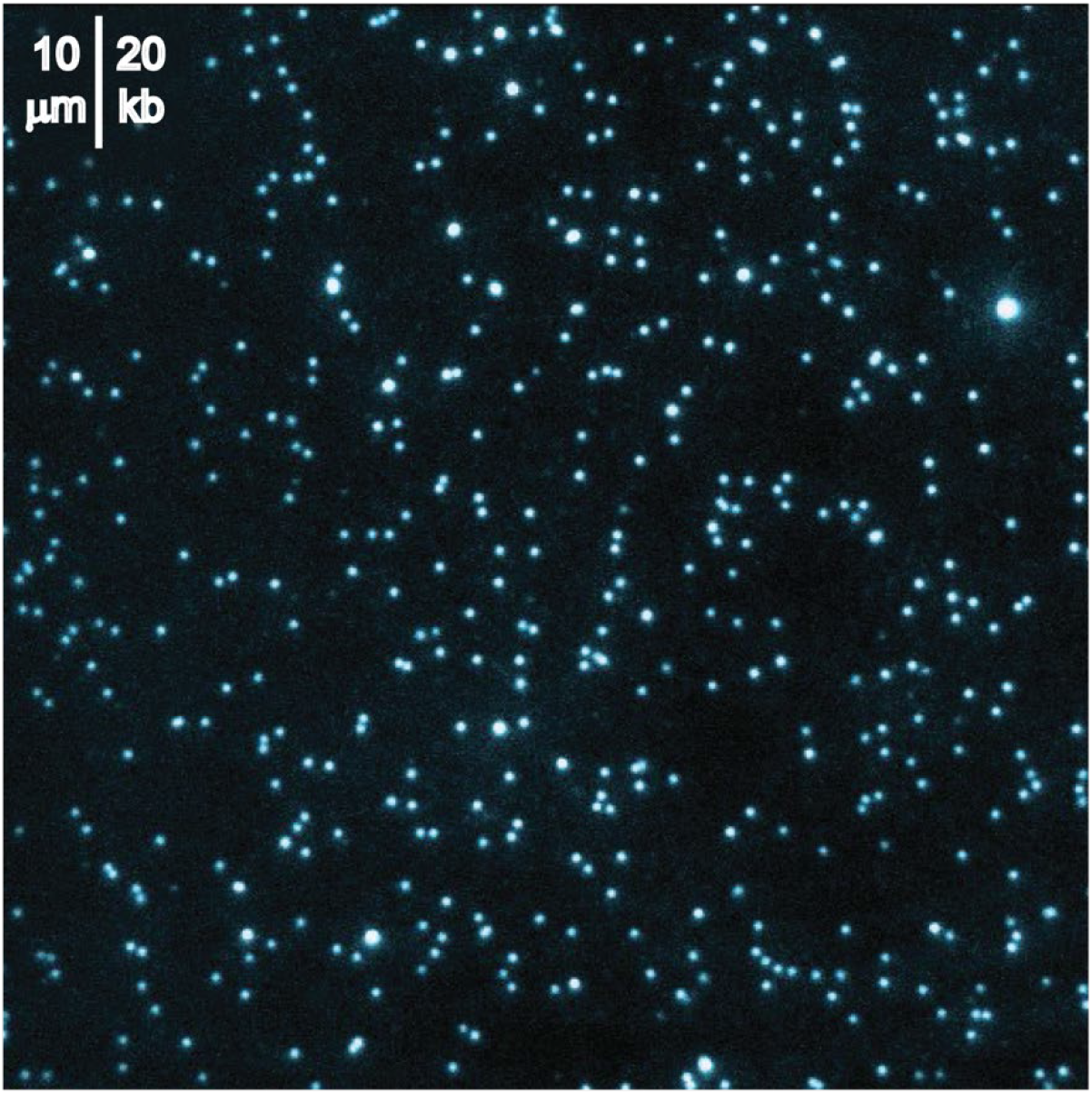
Representative field of view showing the lack of replication products in the leading and lagging-strand synthesis assay after omitting ATP from the DnaB loading phase. The image was recorded 2 min after replication initiation, where if helicase loading was successful, we would expect to see long, replicating DNA products (Fig. 1C and fig. S3) instead of these un-replicated DNA templates.

**Fig. S3.**
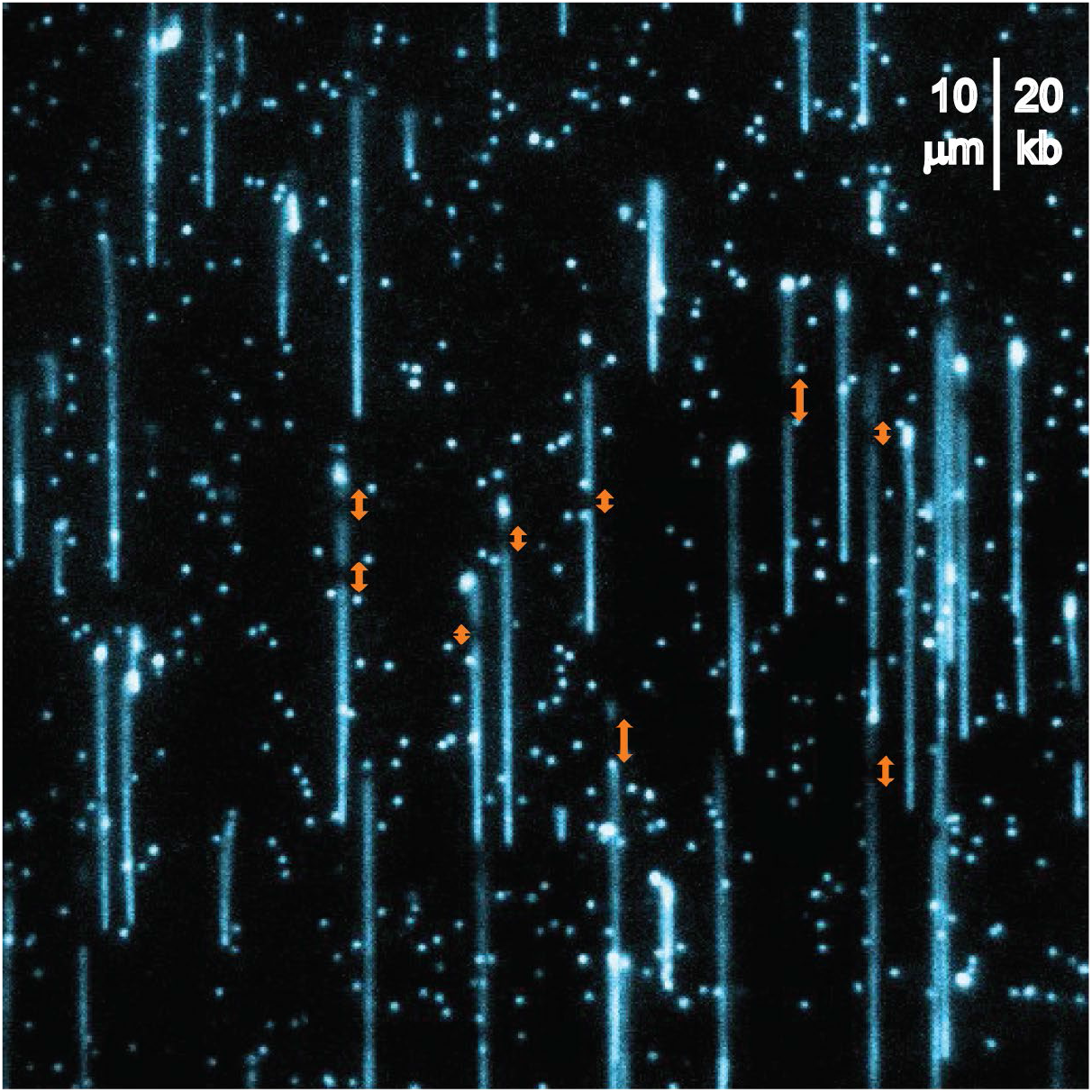
Representative field of view of simultaneous leading- and lagging-strand replication in the absence of ATP (and all other rNTPs). Arrows indicate sporadic ssDNA gaps in the replication products as a result of inefficient priming of lagging strand Okazaki-fragments by DnaG primase using dNTPs.

**Fig. S4.**
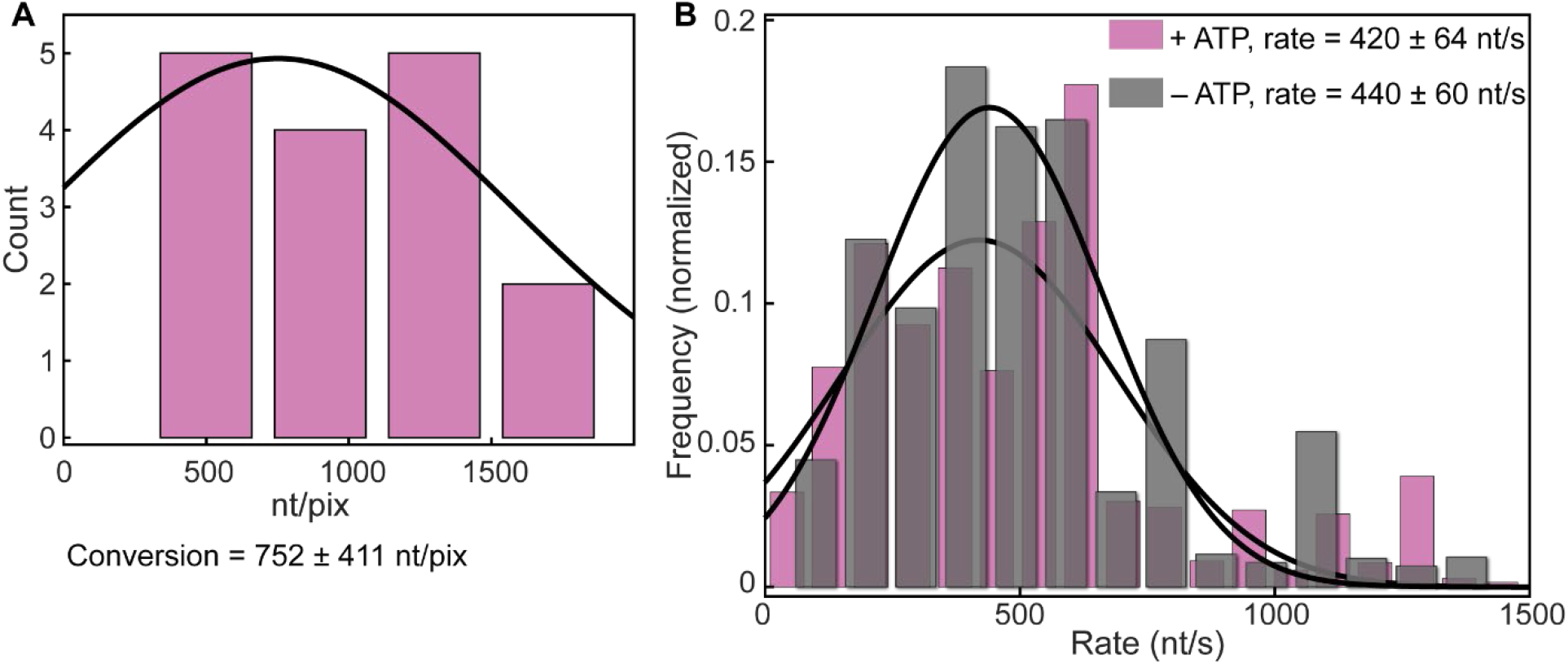
**(A)** Histogram of the number of nucleotides per pixel in leading-strand synthesis. (*N* = 16 molecules) **(B)** Rate histograms of leading-strand replication in the presence (purple, *N* = 80 molecules) and absence (gray, *N* = 116 molecules) of ATP. Errors represent s.e.m.

**Fig. S5.**
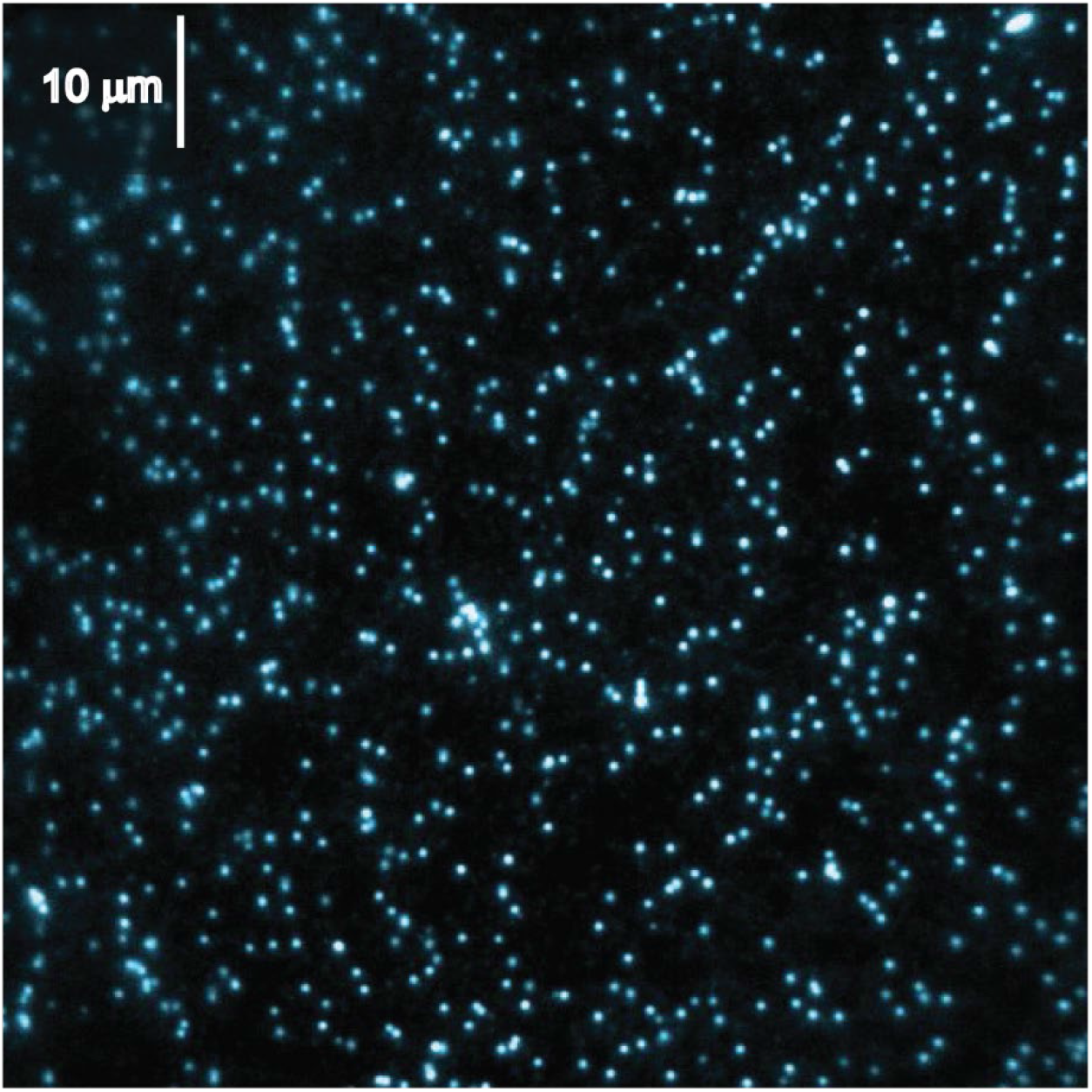
Representative field of view demonstrating the lack of replication when DnaB is omitted from the leading-strand synthesis reaction. The image was taken 2 min after replication initiation. In contrast to Fig. 1C and fig. S3, no replication products were observed in the absence of DnaB. Strand-displacement synthesis is possible by the Pol III holoenzyme and SSB, but only at much higher concentration of dNTPs (*44*).

**Fig. S6.**
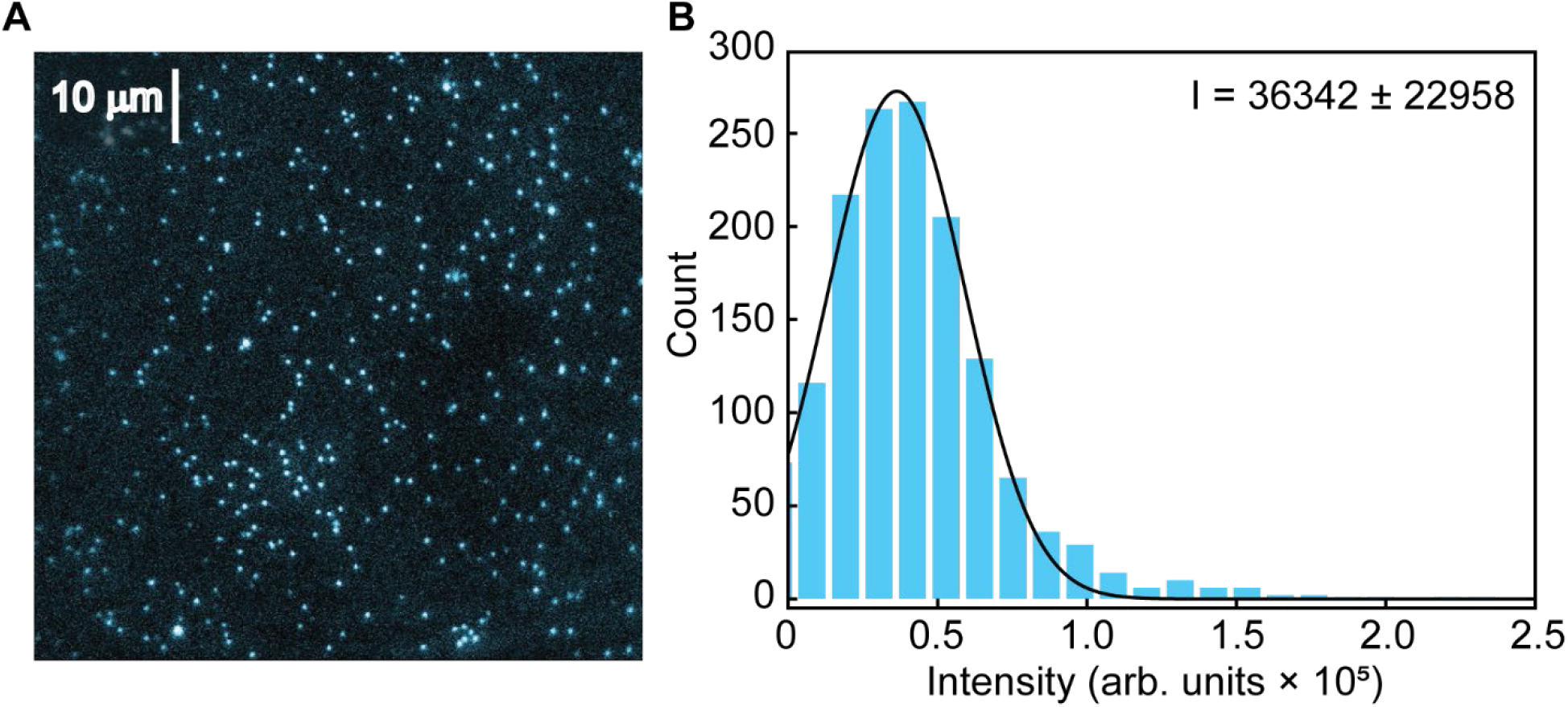
Calibration of ssDNA staining by SYTOX orange. **(A)** Typical field of view showing M13 ssDNA stained with SYTOX. **(B)** Histogram of the intensity of SYTOX-stained M13 ssDNA. The black line represents a Gaussian fit to the data. Using the mean intensity and the known length of M13mp18 (7249 nt), a conversion factor of 5.0 ± 4.5 nt/arb.unit was obtained.

**Fig. S7.**
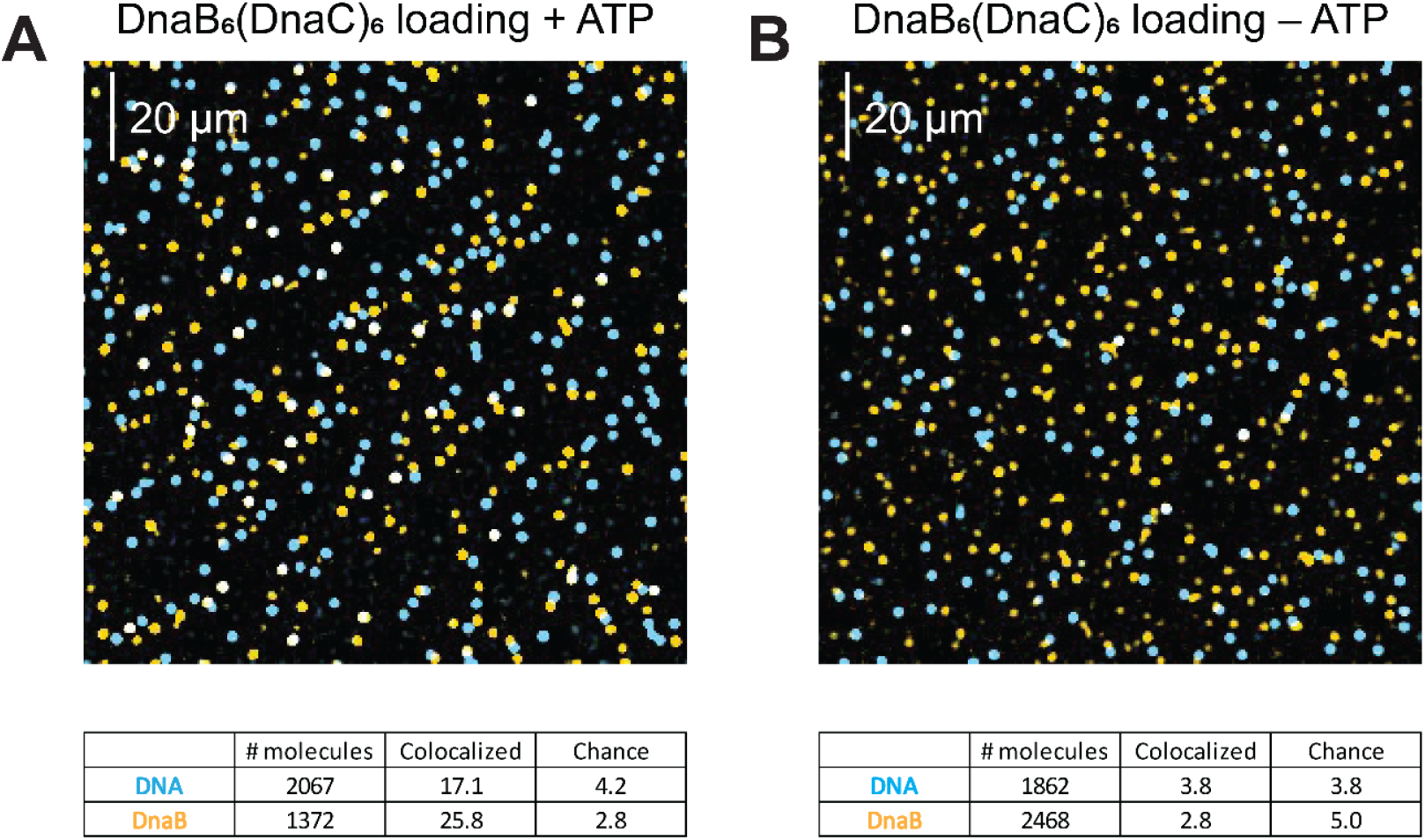
Effect of ATP on DnaB helicase loading. **(A)** Typical field of view of fluorescent DnaB_6_(DnaC)_6_ (orange) loading on the rolling-circle DNA template (blue) in the presence of ATP. Successfully loaded events appear as white. The table shows the total number of molecules analyzed, colocalization (%) and colocalization by chance (%). **(B)** Typical field of view of fluorescent DnaB_6_(DnaC)_6_ (orange) loading in the absence of ATP. Successfully loaded events appear as white. The Table depicts the total number of molecules analyzed, colocalization (%) and colocalization by chance (%).

**Fig. S8.**
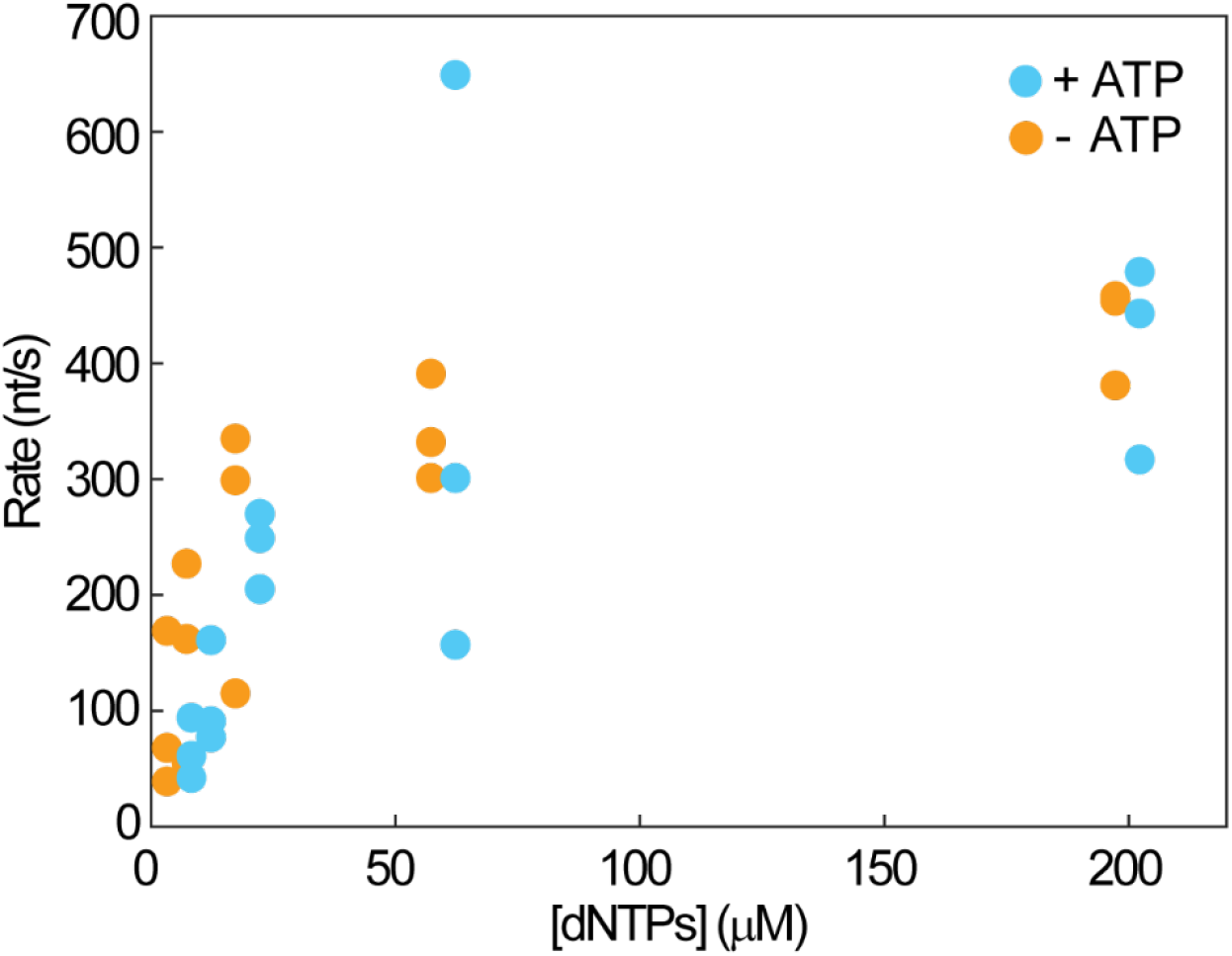
Scatter plots showing replication rates as a function of dNTP concentration in the minimal assay, with (blue) and without (orange) 1 mM ATP. Points represent the mean rate obtained from each replicate experiment (*N* = 3 independent experiments, at each dNTP concentration used; the number of molecules observed ranged between 20 and 100 in each experiment). The [dNTPs] refers to the concentration of each of the four dNTPs.

## Notes

### Competing Interest Statement

The authors have declared no competing interest.

